# Spatial autocorrelation: bane or bonus?

**DOI:** 10.1101/385526

**Authors:** Matt. D. M. Pawley, Brian H. McArdle

## Abstract

Spatial autocorrelation is a general phenomenon within biogeographical studies. However, considerable confusion exists about how to analyse spatially autocorrelated data collected using classical sampling methods (e.g. simple random sampling). We show that the two common discordant views about autocorrelated data depend upon the desired scale of inference. Although inferential statistics seeking to generalise to different unsampled spatial areas need to be adjusted for autocorrelation, if the inference is restricted to the area from which samples have been taken, then standard tests are applicable. In the latter case, incorporating autocorrelation into the model may actually improve the precision and power of the analysis. We found that the scale of spatial inference is rarely discussed, despite being of central importance to any spatial analysis. We suggest that spatial inference should rarely be formally generalised to unsampled areas, since it typically requires some assumption of stationarity, and is thus vulnerable to accusations of “pseudo-replication”.

## Introduction

> *“everything is related to everything else, but near things are more related than distant things”* (Tobler 1970).

Ecologists may argue that they have long been aware of Tobler’s ‘first law of geography’—the phenomenon of spatial autocorrelation (Student 1914, Moran 1950). A belief, still commonly held by ecologists, is that classical statistical methods using design-based inferences (e.g. Simple Random Sampling (*SRS*)), are inappropriate for spatially autocorrelated data. Following the paradigm introduced in Neyman’s prominent paper (1934), the standard argument is that in the presence of spatial autocorrelation, the sampled values are not independent, so you have less information than you think and your estimated degrees of freedom are too large. Consequently, the variance is underestimated and confidence intervals are too small (or the type I error is under-reported) (Cliff and Ord 1981, F. Dormann et al. 2007, Kissling and Carl 2008, Bini et al. 2009, Rousset and Ferdy 2014). Contrary to this opinion, classical sampling theory has been demonstrated as quite applicable to spatial data (de Gruijter and ter Braak 1990, Brus and DeGruijter 1993). These papers elegantly demonstrated that classical statistics used with design-based methods are not only valid for autocorrelated data, but actually require fewer assumptions than model-based (geostatistical) methods that explicitly model the autocorrelation. Although some ecologists have acknowledged this (Legendre 1993), many continue to state that the presence of spatial autocorrelation complicates their analysis, and reduces the real power and precision of statistical tests (Carroll and Pearson 1998, Osborne et al. 2001, Ferguson and Bester 2002, Fisher et al. 2002, Bester et al. 2002, Diniz-Filho et al. 2003, Schwarz et al. 2003, F. Dormann et al. 2007, Beale et al. 2010, Marrot et al. 2015).

How is it possible to have such disparate views about the same effect? Is one of the views incorrect? If both views are valid, when can autocorrelation be ignored and when must it be accounted for? Here, we discuss the mechanism that acts to separate this dichotomy. We will identify those situations in which classical analytical methods are inadequate (and why), and we will show that in those circumstances where classical methods are valid, autocorrelation is actually a bonus, providing an opportunity for ecologists to *improve* the precision and power of their analyses.

### Temporal vs. Spatial Autocorrelation

Misconceptions about classical methods and spatial autocorrelation almost certainly arose because autocorrelation has been widely perceived as a problem with measurements in times series (Chatfield 2004). In fact, autocorrelation in time series data is not a problem when one can apply the same rules of inference commonly used in design-based methods. If a time series analysis only required the parameters describing the time-period spanned by the *sample extent*, i.e. the domain [spatial and/or temporal] within which measurements are recorded (Wiens 1989, Dungan et al. 2002), then all the statistics derived from a random sample from that domain would require no correction for autocorrelation.

For example, consider Fig. 1, which shows a time series of the mean daily humidity from the Aranjuez agrometeorological station (Madrid) between 2004 and 2010. Suppose 20 measurements were randomly taken (red triangles) within 2004 (the sample extent, as depicted by the grey area). This scenario has three different means that may be discussed: (1) the sample mean, 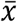, of the 20 sample points; (2) the realisation mean, µ_R_, equivalent to the mean of all data (sampled and unsampled) within the sample extent, and (3) the process mean, µ_p_, which equates to the mean of the stochastic process generating the time series (and is also asymptotically equivalent to the mean of a ‘long-term’ time series. In probability theory, the observed data are often considered to be only one possible outcome that the stochastic process *could* (hypothetically) generate—the possibilities that could happen are known as different ‘realisations’ (Hamilton 1994).

**Figure 1.**
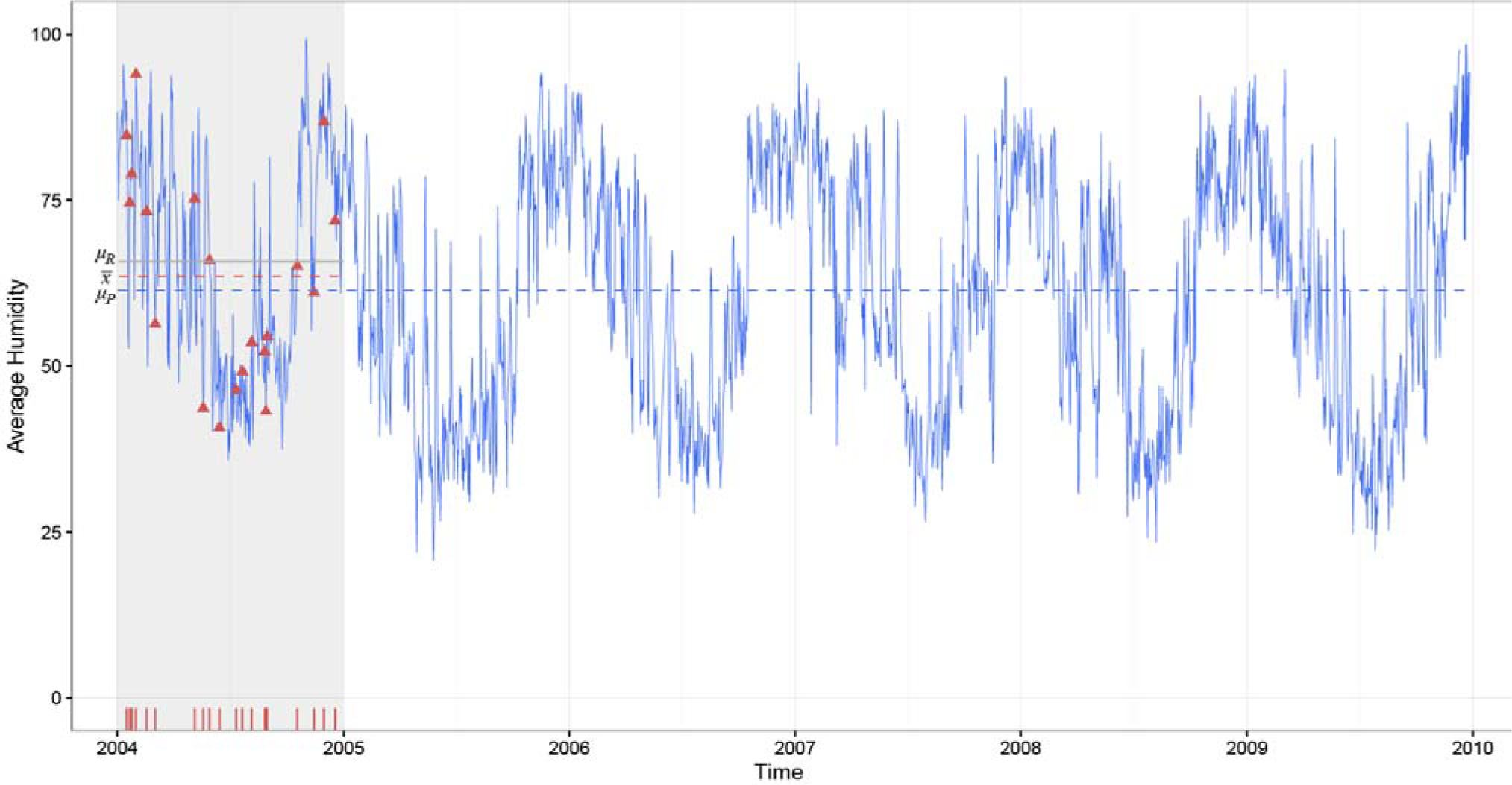
A time series showing average daily humidity at Aranjuez station (Spain) between 2004 and 2010. Twenty random samples (red triangles) were taken within a particular period of interest (the sample extent [shaded in grey]) and used to calculate a sample mean, 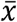 [red line]. The population mean of the sample extent (µ_R_) is depicted by the horizontal grey line, and the mean of the entire time series (µ_p_) is depicted by the dashed blue line.

If the purpose of the analysis in Fig. 1 was to estimate the realisation mean, µ_R_, then the estimate of mean humidity would have confidence intervals (and standard errors) correctly calculated using classical methods that ignore the autocorrelation. That is, if inferences are limited to the period from which we sampled, then autocorrelation is not a problem.

Time series analyses are often used to predict future values (i.e. forecasting), and problems with autocorrelation arise when we wish to generalise our descriptions from the sampled period to other unsampled periods, or to the long term behaviour of the stochastic process (e.g. µ_p_) that generated the sampled year’s records. Such analyses require considerably more restrictive assumptions; in particular, stationarity must be assumed, such that the properties of the system you have measured are the same as those of the spatial region, or period of time, you wish to generalise to [see Myers (1989) for a discussion on how restrictive this assumption must be].

Historically, ecologists have proved well aware of the dangers of attempting to consider a result taken at one place and/or time, and using it to make inferences about other places and other times (i.e. other realisations). Believing the properties of your single field to extend beyond your sampled area so as to generalise to other (unsampled) areas is, at best, optimistic. So in contrast to times series, when sampling an area, the inference space (the space that we wish to make statistical deductions about) is typically restricted to a particular finite area, and the standard error of the sample mean refers to the population contained within the area encompassed by the sample extent, i.e. the realisation mean. If this is the case, then classical statistical theory will perform as intended. SRS, for instance, would yield independent sampling units and an unbiased estimate of the population parameters (e.g. mean and variance of the (finite) population of interest) (de Gruijter and ter Braak 1990).

If you wish to generalise your inferences beyond the population bound by your sample extent, then greater consideration of autocorrelation is required. In the presence of positive autocorrelation, the variability of data within the sample extent is often less than that of the process, leading us to possibly overestimate the precision of our sample mean with respect to the process mean (Pimm and Redfearn 1988, McArdle 1989). In this case, a correction is required for the fact that your particular realisation (the data from which you have sampled) may not describe the set of all possible realisations very well. Generalising inferences beyond the sample extent means that the sample scheme cannot be properly design-based since inference includes unsampled areas. Consequently, if results are then generalised beyond the sample extent, then using a design-based scheme such as SRS to sample a specific (autocorrelated) region will not ensure stochastically independent errors. In this respect stochastic independence depends not only on the sample design and the underlying population autocorrelation structure, but also on the inference space that is chosen. However, the smaller the *range* of autocorrelation (i.e. the distance at which points became independent) relative to the size of your sample extent, the more representative of the total process your realisation becomes. The efficacy of a classical 95% confidence interval using the -distribution (i.e. 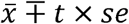) to estimate the realisation and process mean, and the effect of the range of autocorrelation is demonstrated via simulation in the next section.

### A simulation study demonstrating the importance of inferences

Fig. 2 shows two surfaces, each representing a simulated spatial distribution of organisms within the same sized sample extent. These surfaces were created using a positively autocorrelated random field (with a spherical variogram structure) using the GaussRF function within the RandomFields library in the ‘R’ software package (Martin Schlather 2001, R Development Core Team 2015). Since changes in populations are often considered to be the result of birth and death processes, the processes were simulated on a multiplicative scale (Legendre and McArdle 1997). Consequently the variogram (autocorrelation) structure was chosen to represent the spatial distribution of organism density on the log scale. Further stochasticity was introduced by using the (exponentiated) random field values as λ when generating from a Poisson distribution. Surface A was generated with a ‘short-range’ autocorrelation process (the range was around 20% of the surface) and surface B was generated with a ‘long-range’ autocorrelation process (80% of the surface), but all other stochastic parameters used to generate these surfaces were identical.

**Figure 2.**
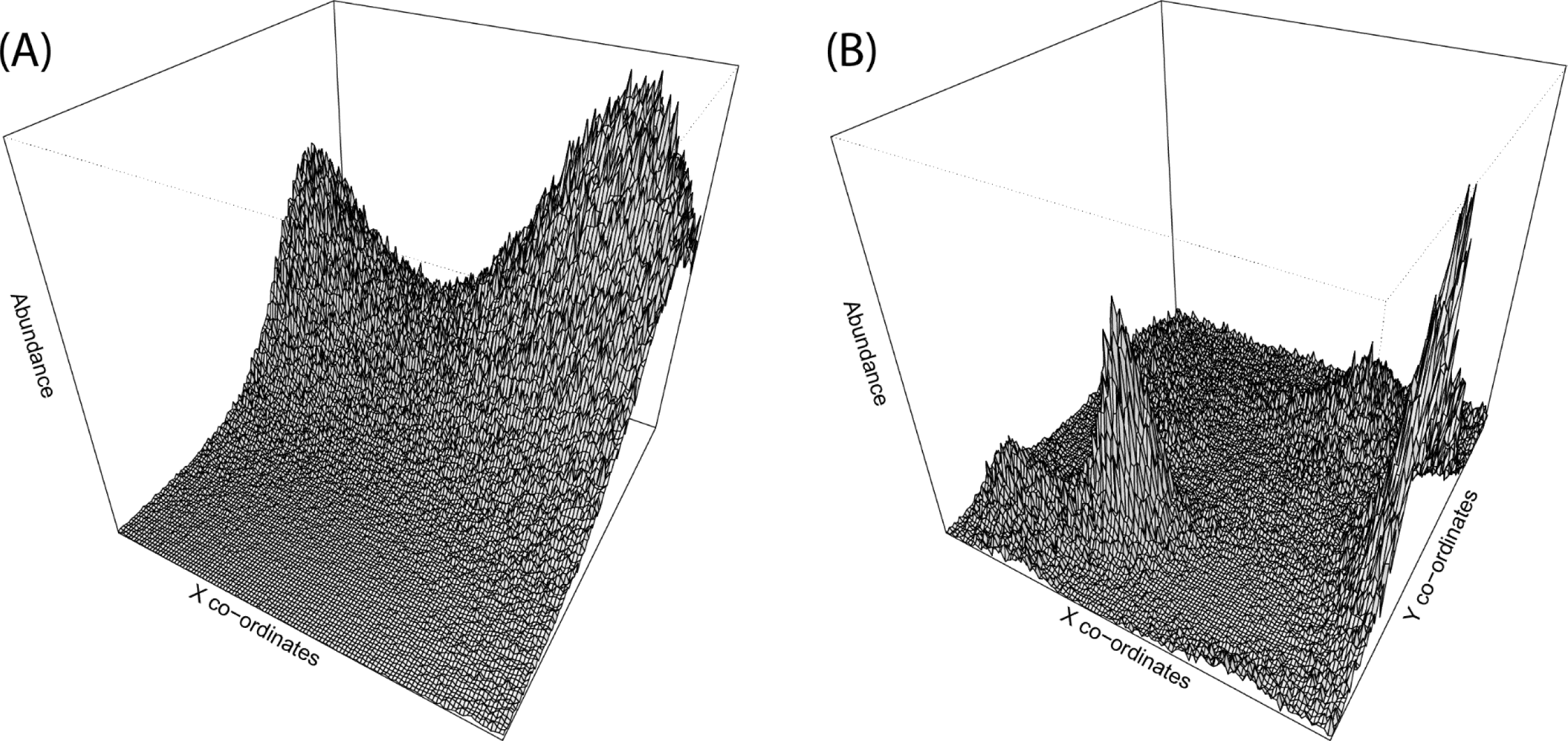
Two equal-sized surfaces simulated to represent different spatial distributions of organisms. The range of autocorrelation (relative to the extent of the surface) is relatively ‘short’ in B and ‘long’ in A.

The distributions shown in Fig. 2 are a single (exemplar) realisation of a simulation which stochastically generated 10,000 surface realisations each of the ‘long-range’ and ‘short-range’ autocorrelation structure. For each realisation, the following statistics were calculated:

1. the realisation population mean (µ_R_);
2. a sample mean 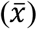 from a single SRS (n = 100);
3. a 95% confidence interval (CI) for the population mean using the classical statistical method (i.e. 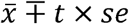).

In addition, the process means (µ_p_) for the ‘long-range’ and ‘short-range’ autocorrelation structure were also calculated. The R-code for the simulation is supplied in Supplemental 1 (S1).

The 95% confidence intervals encompassed their respective realisation means on approximately 94% of samples from ‘short range’ realisations and 91% or ‘long range’ realisations (Table 1)—the stochastic processes generated highly over-dispersed data, so coverage for the realization means were slightly lower than 95%. In contrast, coverage of the process mean was much lower than 95% and depended on the range of autocorrelation (66% and 33% for the short- and long-range surfaces respectively). Confidence intervals from the ‘short range’ realisations (e.g. surface B) had a smaller range of autocorrelation relative to the sample extent, so the sampled data were, on average, more representative of the total process than data from the ‘long range’ process and consequently confidence intervals covered the process mean at a higher rate than the ‘long range’ process. Clearly, if you sample a large area (relative to the range of spatial autocorrelation), then your generalisation is likely to be more reliable than if you only sampled a small one.

| Inference | 'Long range' autocorrelation<br>(e.g. Surface A) | 'Short range' autocorrelation<br>(e.g. Surface B) |
| --- | --- | --- |
| *Realisation mean* | 94.4% | 91.3% |
| *Process mean* | 33.1% | 66.2% |

One commonly overlooked consideration about autocorrelation is that it is not the data values that must be independent, but the residuals after the model has been fitted (ANOVA, regression, t-test etc.) (McArdle 1996). If a hypothesis test refers to parameters describing the process, then independence of errors may be achieved by the inclusion of those variables that help determine the spatial distribution. For example, some gregarious animal might be present at a location because of the presence of the particular habitat type; including some measure of the habitat in the model may remove much of the spatial autocorrelation from the residuals. Inclusion of locational covariates (such as longitude and latitude) can also act as a proxy to some determining covariate variable and help remove spatial autocorrelation from the residuals (Legendre 1990).

### Regression

The classical model-based technique of regression aims to describe the linear dependence of a response variable (*Y*), on an explanatory variable (*X*), correcting for the influence of other *X*’s. If the *X* values used in the estimation procedure are representative of the sample extent (e.g. from a random sample) then the description (or prediction) identified will be valid for the population within the sample extent, irrespective of any underlying autocorrelation. The error variance and inference about the slope parameters (standard errors and tests) will be correctly estimated, but only for statements about the site in question. This may be of limited usefulness.

Most ecologists would like to generalise any apparent relationships between variables to a larger class of sites, or at least be sure that any apparent relationship was not simply an accident of having autocorrelated data. If the range of autocorrelation is large relatively to the area under consideration then it is quite possible to detect a relationship, even if no real relationship would be apparent if a larger extent was studied. The problem arises when a ‘hill’ in the response (*Y*) surface is accidentally aligned with a ‘hill’ (or ‘valley’) in the explanatory variable (*X*) surface (Legendre 1993). The alignment of *X* and may occur due to another variable that is missing in the model (*Z*), which is autocorrelated and related to both *X* and *Y*. The impact of autocorrelation on this *X*, relationship is due to the missing causal variables. In other words, your model was actually incomplete because you left out an important explanatory variable. Ideally, these *Z* variables would be measured and the *X*,*Y* relationship would be clarified by incorporating them into the model (e.g. by multiple regression, or partial correlation). If you include such *Z* variables, the residuals are no longer autocorrelated. As is well known in regression, the validity of the description and inferences about the parameters depends crucially on the adequacy of the model. If the model is wrong then the parameter estimates and the tests upon them may be meaningless (Hawkins 2012). Our ability to generalise is also dependent on the assumptions of the model. In particular, a crucial assumption is that all the predictor variables upon which *Y* depends are in the model; or, if they are not, then the excluded ones are uncorrelated with the included ones (and can therefore be relegated to the error term with impunity).

We are pseudo-replicated, though somewhat informally, if we use the results from one study to suggest trends and hypotheses about other areas. We might accept the generality of the result when many studies in many places and times seem to reproduce it. The formal generalisation of the result beyond our initial study area relies on there being enough studies to merit a meta-analysis (Hunter et al. 1982). Ecologists need to ask: what inferences do we wish to make about our sampled population? If you wish to use a generalised inference and adjust your test according to the relevant autocorrelation, then you will be generalising your inference to encompass all those areas with the same autocorrelation structure. Is this of practical use—will your readers accept the pseudo-replication?

### Exploiting spatial autocorrelation

Consider the following statement:

> *“For [other] applications, such as estimating the mean of an area, however, samples should be spaced more widely than the range* [of autocorrelation]*, to avoid the effects of spatial autocorrelation.”* (Burrough 1995).

Such advice is misleading. In fact, if the sampling scheme positions units at distances much greater than the range of autocorrelation, then the efficiency of the design is actually decreased (towards SRS). As long as inference is restricted to the sample extent, then the adage that ‘sample points are autocorrelated and are therefore not independent’ can actually work in your favour. If the underlying data are autocorrelated, then a sample unit also contains information about its neighbouring points, so you have information about a larger area of the finite population than the simple sample size implies.

Ecologists frequently use methods that exploit the spatial correlation present in field studies, e.g. stratification (for sampling) or restricted randomisation (e.g. blocking, Latin Squares), for experiments. In both cases, areas of similar values are sampled separately from one another, and among-strata variation can be removed from the error term by appropriate modelling methods (e.g., Analysis of Variance). More sophisticated analyses that explicitly model spatially correlated errors are used in agricultural studies to improve precision and power (Brownie and Gumpertz 1997).

Another common method that exploits autocorrelation is systematic sampling. Systematic sampling is widely used in practice, due in large part to its simplicity and convenience, but also because of the intuitive appeal of spreading sampling units evenly over the population, thus ensuring ‘good coverage’ (Singh and Singh 1977, Iachan 1982). Standard use of a random-start, systematic sampling design (SYS) equates to randomly positioning a theoretical grid (with resolution dependent upon the size of the sample) over the region of interest, and measuring the value of the spatially distributed variable at the grid intersections. Provided there are no periodic (waveform) patterns in the population’s surface of values, then, in the presence of positive autocorrelation, SYS is likely to give an estimate of the population mean that is as good or better than any other sampling scheme (Hájek 1959).

## Conclusions

In short, the confusion between the perils of temporal autocorrelation in time series and spatial autocorrelation in spatial samples arises from the respective analyses typically having different objectives. Time series methods are designed to allow extrapolation from a finite sample extent to subsequent unsampled time periods [a *generalised* inference space – analogous to the ‘super-population’ distribution, a term first coined by Cochran (1946)]. In contrast, classical methods, as applied to spatial samples and occasionally time series, usually restrict their description and inference to the sample extent (a *finite* inference space).

If you have autocorrelated data, one of the first steps in determining the appropriate analysis should be examining the scale of inference. This is a step that is rarely mentioned, and commonly disregarded; leading to a relatively common misconception that correct analysis of autocorrelated data automatically requires a more sophisticated technique than the classical methods.

Broadly speaking, there are two types of errors that arise from not determining the appropriate level of inference:

1. The appropriate level inference is to generalise to a larger (possibly infinite) space/population, but the investigator chooses an analysis that is appropriate for a finite region (e.g. ignoring autocorrelation).

This error and its consequences have been well researched and documented.

Ignoring autocorrelation in such circumstances commonly leads to underestimation of the (autocorrelated) parameter variance and confidence intervals/type I error rates that are too small (Cliff and Ord 1973, Griffith 1978, Odland 1988, Dutilleul et al. 1993, Rousset and Ferdy 2014). If inference is to be generalised, there have been a number of possible solutions that have been discussed (see Augustin, Mugglestone, & Buckland, 1998; Dale, Blundon, MacIsaac, & Thomas, 1991; F. Dormann et al., 2007; Pierre Legendre, 1990; Pierre Legendre & Legendre, 2012; Tavare & Altham, 1983).

2. The appropriate level of inference is restricted to a specific surveyed area/finite population, but the investigator chooses an analysis that generalises to a broader inference space (a process method).

The use of some method that incorporates spatial autocorrelation does not necessarily mean that such a method is appropriate for generalisation to a broader inference space. For example, when using model-based geostatistical methods, the standard inference space is restricted to the sample extent. Using a technique designed to generalise inferences will give spuriously large confidence intervals (or decrease the power of the test) and will also give non-optimal parameter estimation by your chosen criterion (e.g. least squares) since you are estimating the parameter of the process rather than the finite population. For instance, incorporating non-independent errors by using, say, a generalised least squares will not only increase the standard error of the results, but will also give a different point estimate from the independent errors model.

In summary, when making inferences around a finite population:

1. The use of SRS means that inference is unaffected by spatial autocorrelation.
2. The use of an efficient sample design (e.g. systematic sampling, stratification) that can exploit any autocorrelation in the data can improve the power of hypothesis tests and decrease confidence intervals.
3. Advantages may also be gained through the use of model-based analyses (e.g. geostatistics) that incorporate the autocorrelation structure of the finite realisation into the inference calculations.

If generalising to a ‘super-population’:

1. Autocorrelation may be problematic depending upon the extent of the generalisation relative to the sampling extent (i.e. how far you want to extrapolate your result). Design-based sampling designs, such as SRS, are no longer applicable—so inference needs to be adjusted for any spatial autocorrelation left in the residuals after fitting the model.
2. The smaller the range of autocorrelation relative to the extent of your realisation (sampling extent), the more representative of the process characteristics your data becomes. Clearly if you sample a relatively large area, your generalisation is likely to be more reliable than if you only sampled a small area.
3. Observational studies can be affected by autocorrelation. Any inferences implying causation would have had to correct for unknown variables (spatial confounding or correlated explanatory variables). The validity of the description and inferences about the parameters depends crucially on the adequacy of the model. If the model is wrong then the parameter estimates and the tests upon them may be meaningless.

It is essential to consider exactly what the generalised population (‘super-population’) is that you wish to make inferences about. We suggest that, in many instances, ecologists should be restricting their inference to the finite population with a definite sample frame. Practical reasons may mean that generalising inference to include non-sampled spatial populations may sometimes be unavoidable. However, it should be kept in mind that such practice means that your inference is open to the accusation of pseudo-replication.

It is worth remembering that the spatial distribution of ecological phenomena is sometimes, in itself, the study goal, not a nuisance. The spatial autocorrelation of ecological phenomena may be a reflection of the underlying processes and functions of interest and spatial statistics provides a toolbox to be able to discover pattern and model ecosystems in a spatially explicit manner.

## Acknowledgements

We would like to thank Adam Smith, Jorge David Aguirre, Libby Liggins and David Eme for their helpful criticisms and suggestions.

